# Quantifying predator functional responses under field conditions reveals interactive effects of temperature and interference with sex and stage

**DOI:** 10.1101/2021.08.15.456391

**Authors:** Kyle E. Coblentz, Amber Squires, Stella Uiterwaal, John P. Delong

**Affiliations:** School of Biological Sciences, University of Nebraska-Lincoln, Lincoln, NE 68588

**Keywords:** feeding rates, foraging, jumping spiders, predator-prey interactions, sexual dimorphism, stage structure

## Abstract

1. Predator functional responses describe predator feeding rates and are a core component of predator-prey theory. Although originally defined as the relationship between predator feeding rates and prey densities, it is now well known that predator functional responses are shaped by a multitude of factors. Unfortunately, how these factors interact with one another remains unclear, as widely used laboratory methods for measuring functional responses are generally logistically constrained to examining a few factors simultaneously. Furthermore, it is also often unclear whether laboratory derived functional responses translate to field conditions.
2. Our goal was to apply an observational approach for measuring functional responses to understand how sex/stage differences, temperature, and predator interference interact to influence the functional response of zebra jumping spiders on midges under natural conditions.
3. We used field feeding surveys of jumping spiders to infer spider functional responses. We applied a Bayesian model averaging approach to estimate differences among sexes and stages of jumping spiders in their feeding rates and their dependencies on midge densities, temperature, and predator interference.
4. We find that females exhibit the steepest functional responses on midges, followed by juveniles, and then males, despite males being larger than juveniles. We also find that sexes and stages differ in the temperature dependence of their space clearance (aka attack) rates. We find little evidence of temperature dependence in females, whereas we find some evidence for an increase in space clearance rate at high temperatures in males and juveniles. Interference effects on feeding rates were asymmetric with little effect of interference on male feeding rates, and effects of interference on females and juveniles depending on the stage/sex from which the interference originates.
5. Our results illustrate the multidimensional nature of functional responses in natural settings and reveal how factors influencing functional responses can interact with one another through behavior and morphology. Further studies investigating the influence of multiple mechanisms on predator functional responses under field conditions will provide an increased understanding of the drivers of predator-prey interaction strengths and their consequences for communities and ecosystems.

## Introduction

Functional responses describe predator feeding rates on prey and are central to understanding predator-prey interactions and the structure of food webs. Functional responses were originally defined as the relationship between predator feeding rates and the densities of their prey (Holling, 1959; Solomon, 1949). In the intervening decades, however, we have discovered a multitude of factors other than resource densities that influence predator feeding rates such as: predator density/interference (DeLong & Vasseur, 2011; Hassell & Varley, 1969; Novak & Stouffer, 2021), temperature (Thompson, 1978; Uiterwaal & DeLong, 2020), predator and prey body sizes (Rall et al., 2012; Vucic-Pestic et al., 2010), habitat complexity (Mocq et al., 2021; Toscano & Griffen, 2013), alternative prey availability (Hossie et al., 2021; Murdoch, 1969), and others. We also have learned that the effects of these factors on predator feeding rates have important ramifications for population dynamics and their stability (Amarasekare, 2015; Beddington, 1975; Coblentz & DeLong, 2020; Murdoch & Oaten, 1975; O’Connor et al., 2011; Otto et al., 2007; Uszko et al., 2017) and are important for understanding how global changes in climate, habitat, and the movement of species are likely to influence predator-prey interactions and communities (Dick et al., 2014; Gilbert et al., 2014; Mocq et al., 2021).

Although we know that functional responses are shaped by a variety of factors, our knowledge is limited in how these factors interact to jointly determine predator feeding rates (DeLong, 2021). One reason for this is that nearly all predator functional responses have been measured under laboratory conditions (Uiterwaal et al., 2018). This creates two important issues. First, generally only a few factors that influence functional responses can be considered simultaneously because adding factors greatly increases the number of experimental units required to measure the functional response. Second, it is often unclear how relevant the treatments used in laboratory experiments are to the variation that organisms experience naturally (Griffen, 2021). For example, studies on how functional responses are influenced by temperature rarely mention how or whether the temperature range used in the experiment corresponds to the actual temperatures experienced by the species in the field. Together, these two issues hinder our understanding of how multiple factors interact to shape functional responses and diminish the relevance of experiments to our understanding of natural populations.

A promising alternative to laboratory functional response experiments are observational approaches to measuring functional responses (Baudrot et al., 2016; Beardsell et al., 2021; Novak et al., 2017; Novak & Wootton, 2008; Preston et al., 2018; Smout et al., 2010). These observational approaches allow functional responses to be measured under natural conditions. This allows for the examination of multiple factors influencing functional responses by taking advantage of natural variation in those factors. Furthermore, this also guarantees that the range of variation in the factors are relevant to those experienced by the organisms in the field.

Here we use an observational approach to examine how temperature, predator interference, and sex/stage simultaneously influence the functional response of zebra jumping spiders (*Salticus Scenicus*) foraging on midges (Chironomidae spp.). Our results show that all three factors interact in a field setting and are the first to demonstrate the temperature dependence of functional responses and asymmetric interference across stages and sex in a natural setting. Together, our results demonstrate the multi-dimensional nature of functional responses and the need to consider this complexity to better understand predator-prey interactions and food webs.

## Materials and Methods

### Study System

Zebra jumping spiders have a holarctic distribution and are common denizens of artificial structures. At our field site at Cedar Point Biological Station, Ogallala, Nebraska, USA (41.2N, 101.6W), zebra jumping spiders are common on the outer walls of buildings. The spiders forage on a variety of invertebrate prey but most of their diet in the summer consists of midges emerging from nearby Lake Ogallala. We therefore focus on their functional responses on these midges (See Results). Zebra jumping spiders exhibit marked sexual dimorphism as adults. Males have enlarged chelicerae and are overall darker and smaller than females (Figure 1A,B). Juveniles exhibit a phenotype similar to the adult females but are smaller and generally lighter in color than females (Figure 1A,C). This species is well-suited for the use of observational approaches as they are readily observable feeding on their prey after capturing it for several minutes to a few hours and the times for which the feeding events are detectible are readily measurable (see Detection Time Surveys below).

**Figure 1.**
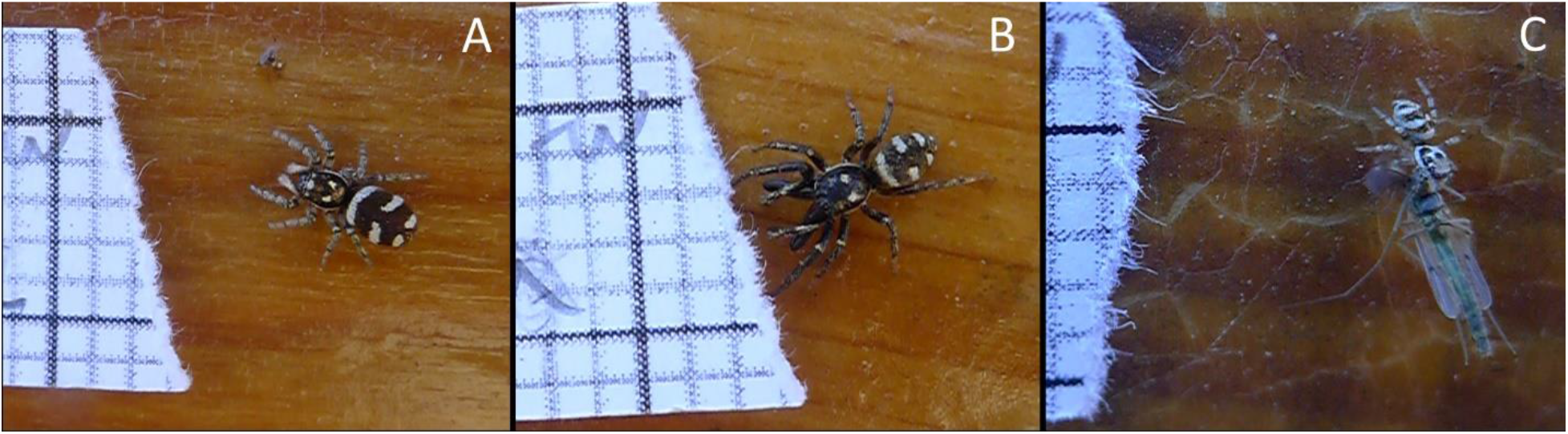
Female (A), male (B), and juvenile (C) zebra jumping spiders. The light gray grids on the grid paper are 2mm.

### Observational Approach to Measuring Functional Responses

We first introduce the observational approach we used to quantify zebra jumping spider functional responses generally and then describe the methods used to collect the requisite data to use the approach and the statistical methods used to fit the functional response models.

The observational approach to estimating predator functional responses relies on the fact that a predators’ feeding rate together with the time over which interactions are detectable gives the proportion of expected time that individuals are observable feeding (Novak et al., 2017; Novak & Wootton, 2008). More concretely, for a species with feeding rate, *f*, the number of prey eaten by that predator over time, *T*, is *fT*. If *d* is the time over which the interaction is detectable (in our case, the time that spiders spend feeding on their prey), the total time a predator is observable feeding is *fdT* and the proportion of time a predator is observable feeding is *fd*. Assuming that individuals have the same feeding rate, then, in a snapshot survey across individuals, the proportion of individuals feeding, *p*, should also be *fd*.

The fact that the feeding rate of a predator can be linked to the proportion of predators observable feeding allows us to connect the two to infer predator functional responses. With only one prey species, individuals are either feeding or not feeding. In this case, we can model the proportion of feeding events as following a binomial distribution. With several surveys of the number of spiders feeding and not feeding across conditions likely to influence predator feeding rates (e.g. prey densities), we can model the number of individuals feeding in each survey *i*, *y_i_*, as

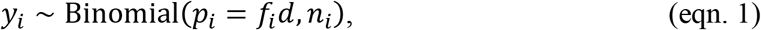

where *p_i_* is the estimated proportion of individuals in survey *i* feeding, *f_i_* is the estimated feeding rate of predators in survey *i*, *d* is the time over which a feeding event is observable, and *n_j_* is the total number of predators in survey *i*. Because the predator functional response describes predator feeding rates across different conditions, by substituting a functional response model for *f_i_*, we can estimate the parameters of that functional response using maximum likelihood or Bayesian methods.

To be able to use this approach, we need several pieces of information required in equation 1. First, from observational surveys, we need the number of predators feeding and not feeding and any associated information to be used in the functional response such as prey/predator densities, temperature, etc. during the survey. We also need an estimate of the detection time *d* or how long, on average, the predators in a survey are observable feeding on prey. Below, we first describe how we performed our feeding surveys. We then describe how we estimated the detection times of zebra jumping spider sexes and stages on midges. Lastly, we describe the statistical methods we used to combine this data to estimate sex- and stage-specific functional responses using the observational method.

### Feeding Surveys

Between May 29, 2020 and June 14, 2020, we performed 155 feeding surveys across 17 building/wall combinations Cedar Point Biological Station. We performed the surveys between 830 and 1600, as the spiders generally were not foraging outside of this time range. We also surveyed specific building-wall combinations at most three times in one day and, if more than one survey was done, we assured that the time between surveys was long enough that feeding events were independent.

Before each survey, we measured the temperature at four to twelve spots along the wall using an infrared thermometer (Raytek Raynger ST, Fluke Corporation, Everett, Washington, USA). We then started at one end of the wall and systematically moved from one end of the wall to the other searching vertically to a standardized height of 1.75 meters. As we moved along the wall, we gave each observed spider a unique ID. We wrote this ID on a piece paper with either a 2 or 6.35mm grid. We then recorded the spider’s sex or stage, whether the spider was feeding, what the spider was feeding on, and, if the spider was feeding on a midge, a description of its size (small, medium, or large). We then photographed the spider with the grid paper visible in the photograph. As we moved along the walls, we also recorded each midge we came across and classified them based on size as for the midges being fed on. From these data, we were able to get the number of spiders of each sex or stage feeding and not feeding on midges, the total number of midges, and an average temperature of the wall during the survey.

After the surveys, we measured the lengths of spiders from the photographs using ImageJ (Schneider et al., 2012). We were unable to get sizes for every individual because some spiders hid or leapt from the wall before they could be photographed, some photographs were not of high enough quality, and one survey was missing photographs. For these spiders, we estimated their size as the mean size for that sex/stage across the experiment.

### Detection Time Estimates

To use the observational approach, one needs an estimate of how long predator feeding events are observable (i.e., detection times). To get an estimates of midge detection times, we performed surveys in which we fed midges to spiders and recorded the length of time from when the spider attacked the midge until the spider subsequently dropped the midge. We performed the surveys between May 29, 2020 and June 13, 2020 and between June 15, 2021 and June 18, 2021. We performed the additional surveys in 2021 after realizing that the detection times surveys performed in 2020 did not cover the full temperature range that occurred during the 2020 feeding surveys. During each survey, we captured midges in clear plastic vials and then placed the vial opening over a spider on a wall until the spider attacked the midge. Spiders were generally returned to the wall after attacking the midge, but, in cases in which the spider refused to return to the wall, they were left to feed inside the vial placed near the wall. We then recorded the attack time, the temperature using an infrared thermometer, and the time when the spider dropped the midge. During the feeding surveys, we also occasionally observed a spider as it caught a midge. When this was the case, we recorded the time, temperature, and the time at which the spider dropped the midge. For all detection time observations, we also took photographs of the spider with grid paper containing a unique ID from which we later measured the spider length in ImageJ as for the feeding surveys. In total, we made 82 observations on females, 18 observations on males, and 17 observations on juveniles.

From this data, we were able to estimate the relationship between midge size, predator size, temperature, and detection times using multiple linear regression through the ‘brms’ package (Bürkner, 2017) in R (v. 4.0.5; R Core Team, 2020). We fit this model after log transforming the detection times, spider length, and temperature to meet model assumptions. We used the default priors with four Hamiltonian Monte Carlo chains with 1,000 sampling iterations and a warm-up of 1,000 iterations. We did not include sex/stage in the model as including sex/stage reduced the predictive ability of the model according to the Widely Applicable Information Criterion (WAIC, Watanabe, 2013).

Using the model fit to the detection time survey data, we estimated an average detection time of zebra jumping spiders feeding on midges for females, males, and juveniles in each survey for which they were present. Partway through the feeding surveys, we standardized observers’ definitions of midge size. For each survey-sex/stage combination, we first determined observer-corrected midge densities of each size for surveys occurring before June 6, 2020. On this date, the observers met and standardized definitions for small, medium, and large midges. Correction factors for each observer were calculated by determining the differences in the number of small, medium, and large midges in surveys post June 6 between each observer’s pre-June 6 definitions of prey sizes and post-June 6 definition of prey sizes. We also used differences in prey sizes from the reclassification of prey sizes from photographs of feeding spiders pre-June 6. The differences in proportions of midges in each size class pre- and post-June 6 were used to correct the pre-June 6 number of midges in each size class using the standardized definition of midge size for each observer. We then calculated the mean spider length in each survey-sex/stage combination and used the results of the regression model to calculate an average detection time incorporating the temperature during the survey and accounting for the sizes of midges available.

### Estimating Functional Responses

To estimate predator functional responses within each stage/sex, we used a model comparison approach by fitting various functional response models to the data and then comparing their ability to predict the data using the Widely Applicable Information Criterion (WAIC, Watanabe, 2013). With observations of the numbers of spiders of each sex/stage feeding and not feeding in each survey, the number of midges present in each survey, the temperature during each survey, and the detection times for each stage/sex feeding on midges, we used equation 1 to fit a suite of functional response models for each stage/sex. We started with a full model for the functional response that included temperature dependence of the predator space clearance rate (*a*) and separate interference parameters for each sex/stage. We modeled temperature dependence of the space clearance rate by assuming that the space clearance rate had an exponential relationship with temperature that could be quadratic (Englund et al., 2011):

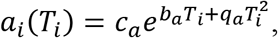

where *a_i_* is the space clearance rate of the predators in survey *i* at temperature *T_i_*, *c_a_*, *b_a_*, and *q_a_* are parameters describing the relationship between space clearance rates and temperature across surveys. With separate interference parameters for each stage/sex, this gave the following full functional response model for the feeding rate in survey *i*, *f_i_*:

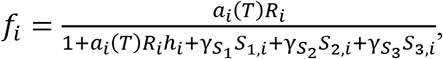

where *R_i_* is the density of midges in survey *i*, *h_i_*, is the handling time of spiders on midges in survey *i*, *γs_j_* are the interference rates of stage/sex *j*, and *S_j,i_* are the densities of stage/sex *j* in survey *i* (Note that the density of the focal sex/stage was calculated as the abundance of that sex/stage – 1 divided by the area of the wall as individuals cannot interfere with themselves). As a simplifying assumption, we assumed that the handling times for each survey were equivalent to the detection times (*h_i_* = *d_i_*). We justify this assumption as the time spent feeding on midges serves as a rate-limiting step for the actively foraging spiders we observed on the walls during the feeding surveys (Novak et al., 2017). In addition to this full model, we also considered the following six models: 1) temperature-dependent space clearance rates and a single, combined interference parameter across sexes/stage, 2) temperature-dependent space clearance rates and no interference, 3) a temperature-independent space clearance rate and separate interference parameters, 4) a temperature-independent space clearance rate and combined interference, 5) a temperature-independent space clearance rate and no interference (Holling Type II functional response), and 6) a null model in which there is a single, across survey average feeding rate and individual surveys only differ from this feeding rate due to differences in detection times.

We fit each of the functional response models for each sex/stage in a Bayesian framework using the program Stan through R using the R package ‘rstan’. Each functional response model was fit using four Hamiltonian Monte Carlo chains with 1,000 sampling iterations following 1,000 warm-up iterations. For the models with temperature-dependent space clearance rates, *c_a_* was restricted to be positive. For the models with interference parameters, all interference parameters were restricted to be positive. For the models with temperature-independent space clearance rates, the space clearance rate was restricted to be positive. Weakly informative or regularizing priors were placed on each of the parameters. For *c_a_* and the temperature-independent space clearance rate *a* we used a Normal(mean = 10, standard deviation = 15) prior derived from invertebrate predators feeding on invertebrate prey in the FoRAGE (Functional Responses Around the Globe in all Ecosystems) database (Uiterwaal et al., 2018). The same prior was also used for the feeding rate in the null model. For *b_a_* and *q_a_*, we used Normal(mean = 0, standard deviation = 1) priors. For the interference parameters, we used Half-Normal(mean = 0, standard deviation = 5) priors derived from the Beddington-DeAngelis functional response fits from Novak and Stouffer (2021).

After fitting each of the functional response models, we used the R package ‘loo’ (Vehtari et al., 2020) to calculate WAIC values for each model, their differences, and the standard error of their differences. We also calculated model weights for each model using the WAIC values (McElreath, 2016). Using these values, we determined the relative abilities of the models to predict the data. To examine the predicted effects of prey densities, interference on predator feeding rates and temperature on space clearance rates, we used the WAIC model weights to perform Bayesian model averaging and produce predictions (McElreath, 2016).

All data and code associated with the analyses are available (see Data Availability Statement).

## Results

### Detection/Handling Times

Detection/handling times decreased with increasing spider length and temperature and decreasing prey size (Figure 2). We estimate that a 10% increase in spider length reduces detection/handling times by 6.6% (95% Credible Interval (CrI) 4.4-9%) and a 10% increase in temperature reduces detection/handling times by 6.2% (95% CrI 1.2-10.9%; Figure 2A,C). Feeding on a medium sized midge reduces the geometric mean detection/handling time by 53% (95% CrI 40-64%) relative to large midges and feeding on small midges reduces the geometric mean of the detection/handling time by 74.6% (95% CrI 67-80%) relative to large midges (Figure 2B).

**Figure 2.**
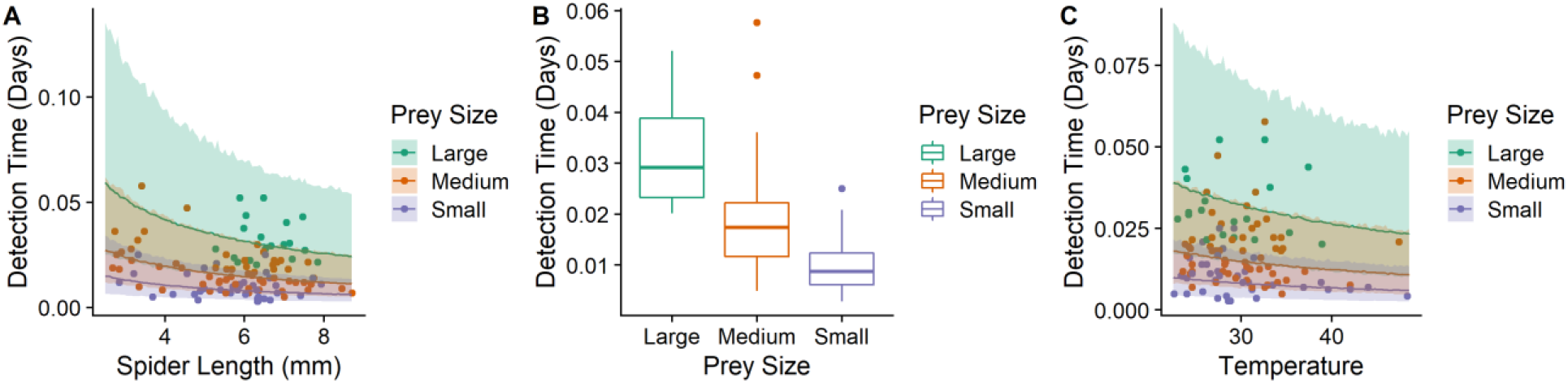
Zebra jumping spider detection/handling times decreased with increasing spider length (A), decreasing prey size (B), and increasing temperature (C).

### Functional Response Estimates

Overall, we used 147 feeding surveys to estimate functional responses for females, 112 feeding surveys for males, and 96 feeding surveys for juveniles. These surveys contained 644 observations of females (average of 4.3 per survey), 286 observations of males (average of 2.5 per survey), and 175 observations of juveniles (average of 1.8 per survey). For females, 41% of the observations were feeding observations of which 92% were on midges, for males, 14% of the observations were feeding observations of which 96% were on midges, and for juveniles, 24% of the observations were feeding observations of which 97% were on midges.

The functional response models considered were ranked differently among sexes and stages (Table 1; See Supplementary Table S1.1-3 for a table including model parameter estimates). For females, the three highest ranked models were the models without temperature dependence of the space clearance rates and made up 92% of the model weight. Models including interference (both combined and separate across stages/sexes) made up 86% of the total model weight. For males, the highest ranked model was the model with no temperature dependence of the space clearance rate and no interference making up 57% of the total model weight. In total, models without interference made up 75% of the model weight. Models with temperature-dependent space clearance rates made up 25% of the model weight. The null model also made up 3% of the model weight. For juveniles, the top four ranked models included interference and made up 81% of the model weight. Models without temperature dependence of space clearance rates made up 56% of the model weight, whereas the models with temperature dependence made up the remaining 44% of the model weight.

**Table 1.**
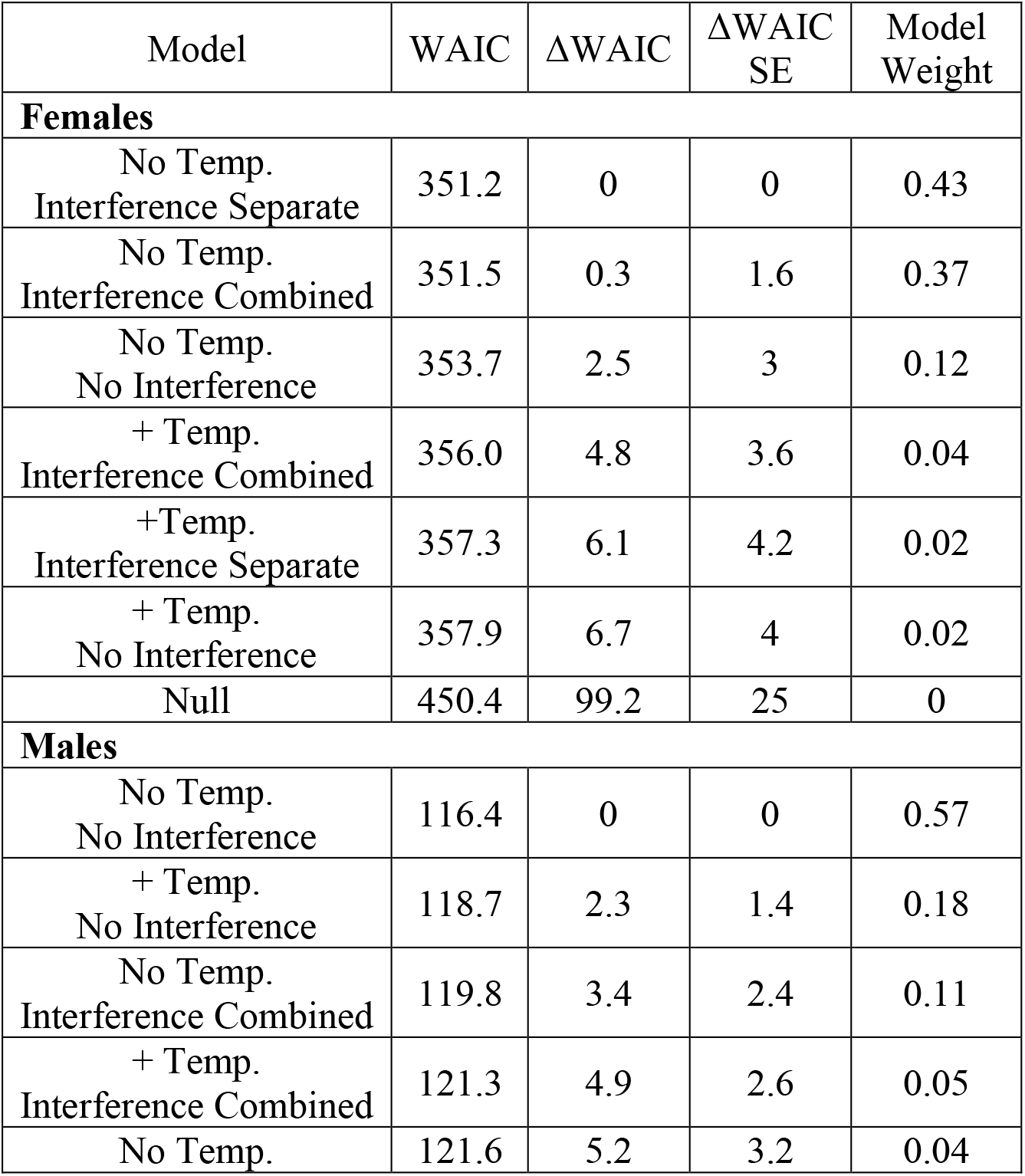
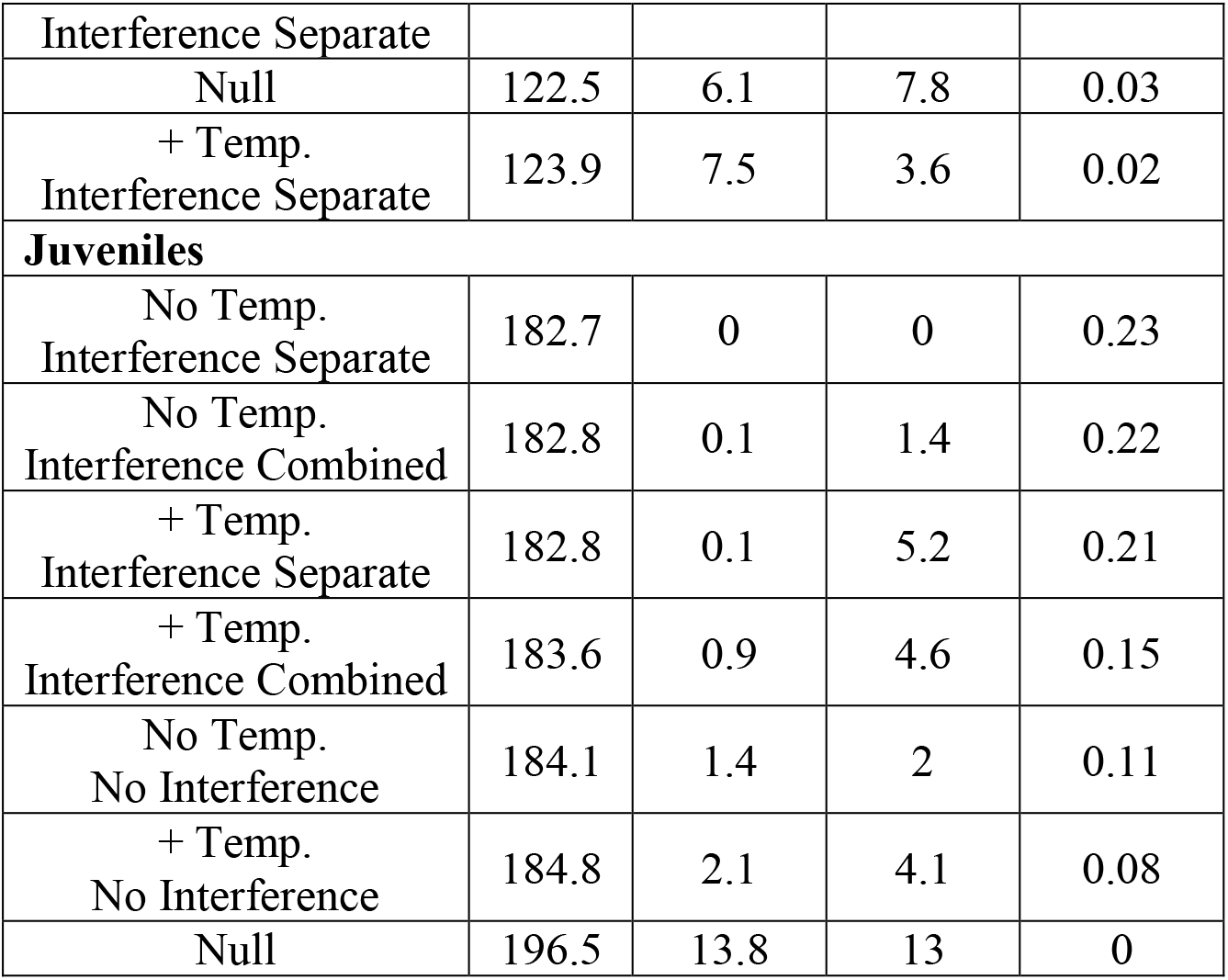
WAIC values for each of the functional response models fit for each sex/stage, Δ WAIC (the differences between the lowest WAIC value and the WAIC value for each model), the standard error of the differences between WAIC values, and model weights calculated from the WAIC values.

Model-averaged predictions of the relationship between midge densities and predator feeding rates show that females exhibit the highest feeding rates across midge densities, with lower feeding rates for juveniles, and the lowest feeding rates for males (Figure 3). The temperature dependence of space clearance rates differed among the sexes/stages (Figure 4). Model-averaged predictions of the relationship between space clearance rates and temperature for females are generally flat, with some weak evidence of a unimodal relationship. Model-averaged predictions for males and juveniles show some evidence of an increase in space clearance rates at high temperatures. The effects of interference also differed among sexes and stages (Figure 5). For males, interference has little effect on feeding rates with similar effects from each sex/stage. Females show reductions in feeding rates from all three sexes/stages with some evidence for greater interference from males and juveniles. Juveniles also show reductions in feeding rates with densities of all three sexes/stages with some evidence of greater interference from females and males.

**Figure 3.**
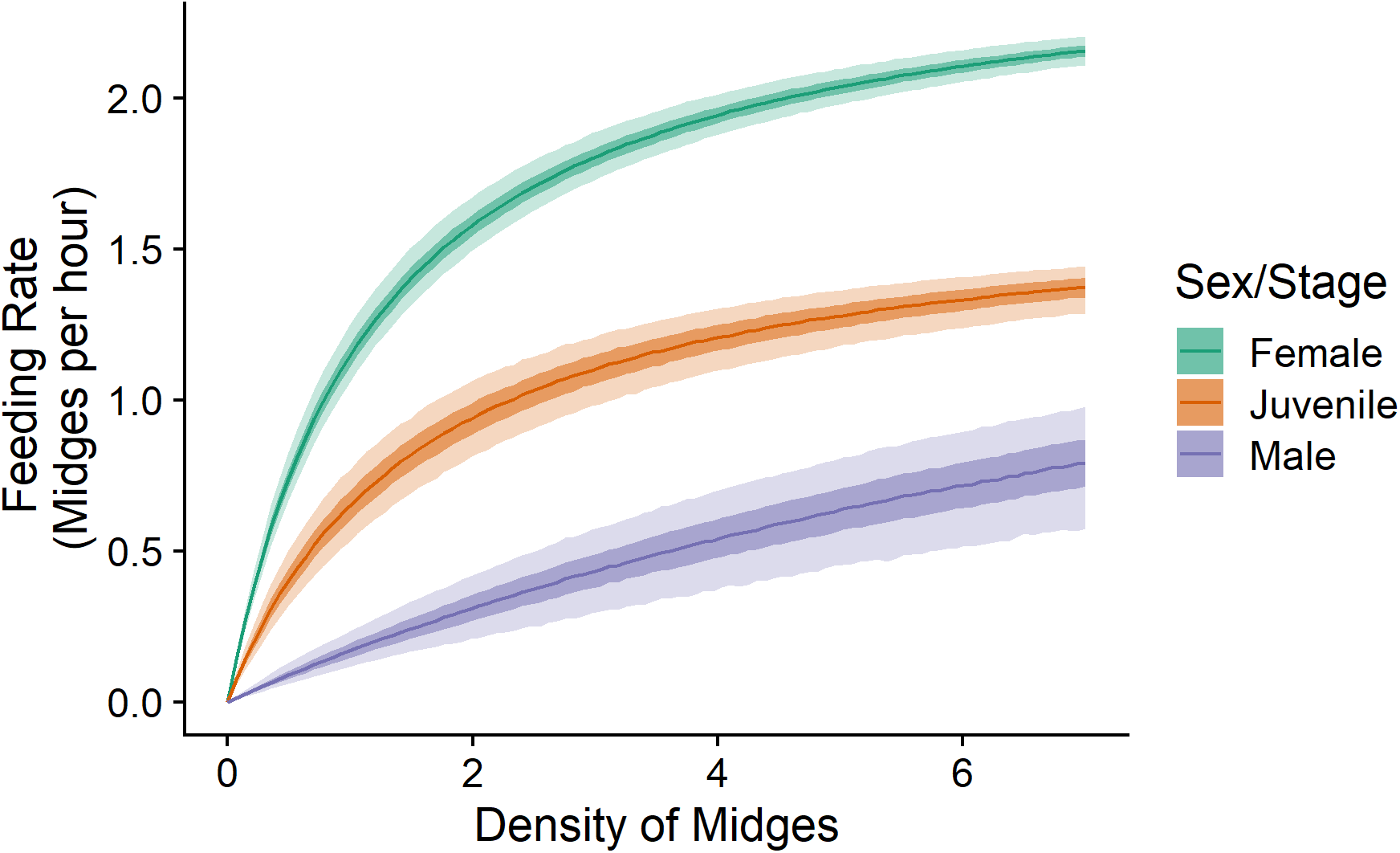
Model-averaged predicted functional responses of female, juvenile, and male zebra jumping spiders. The lines represent predicted median feeding rates for each sex/stage evaluated at the across-survey means for each variable other than midge density. The lightest ribbon around each line represents the 90% prediction interval and the darker ribbon represents the 50% prediction interval.

**Figure 4.**
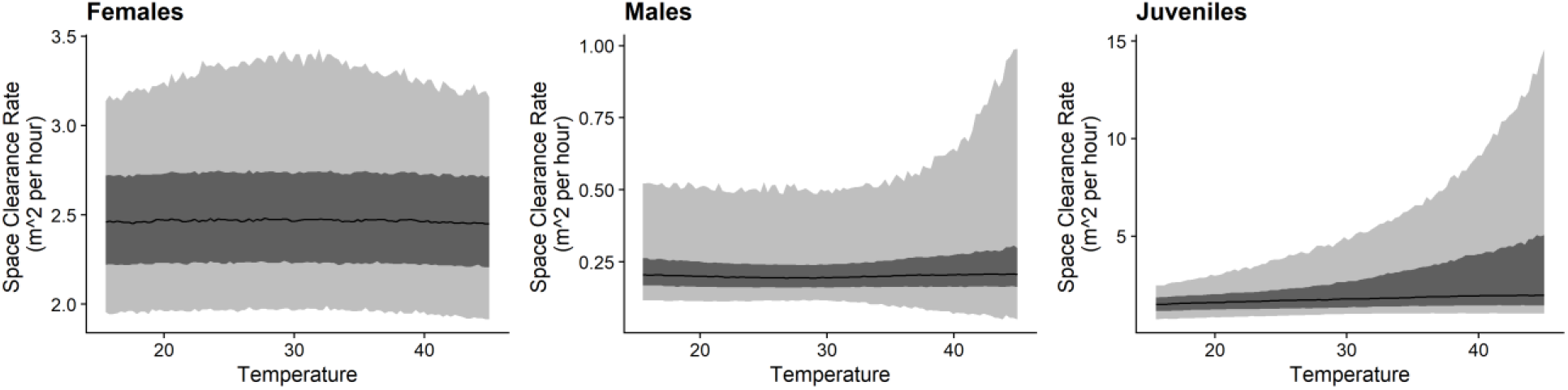
Model-averaged predictions of the relationship between temperature and space clearance rates differ among female, male, and juvenile zebra jumping spiders. All sexes and stages generally show little effect of temperature with some evidence of a unimodal relationship in females, and some evidence of increases in space clearance rates at high temperatures in males and juveniles. The black line in each panel is the median prediction, the dark ribbon represents the 50% prediction interval, and the light ribbon represents the 90% prediction interval.

**Figure 5.**
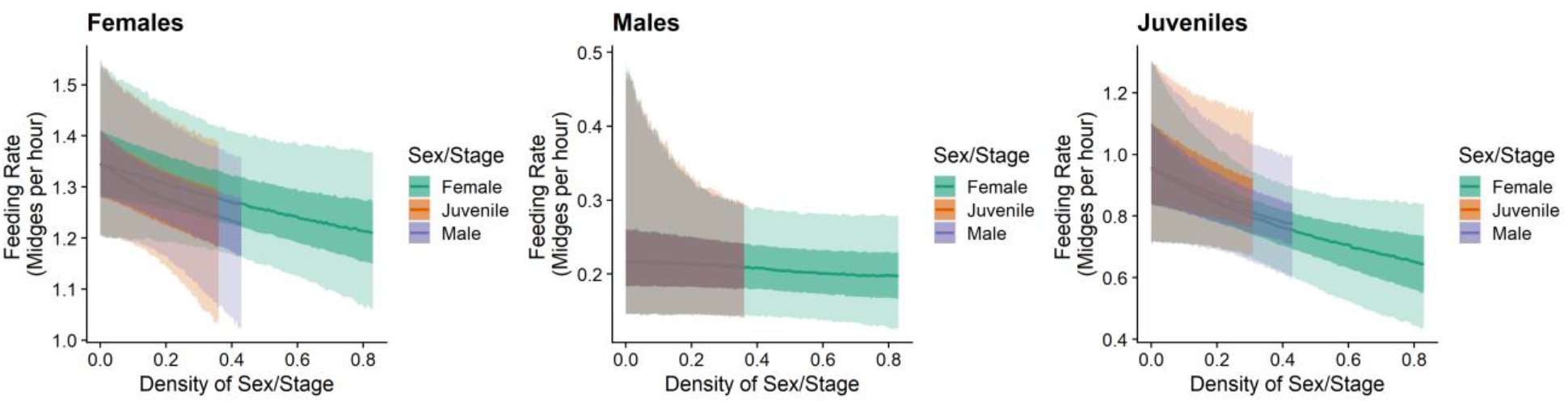
Model-averaged predictions of the effects of interference on zebra jumping spider feeding rates differ among sexes and stages. Each panel shows the effect of the density of each stage/sex on the focal stage/sex’s feeding rate across the observed range of densities in the surveys assuming the densities of the stage/sexes are zero. Prey densities and temperature are at their means across the surveys.

## Discussion

Understanding the factors that influence predator feeding rates and, hence, their functional responses, is central for understanding predator-prey interactions and their consequences. However, studies measuring the effects of multiple factors on functional responses under field conditions remain rare. Here we show that not only do sexes and stages of zebra jumping spiders differ in the relationship between feeding rates and prey densities, but they also differ in how those functional responses are shaped by temperature and interference.

In general, our study predicts that, under similar conditions, females will exhibit the steepest functional responses followed by juveniles and then males. This is despite males being generally larger than juveniles (mean total length of males = 5.01mm, SD = 0.7; mean total length of females = 3.78mm, SD = 0.66), and functional responses generally being expected to increase with body size at least over some range (Rall et al., 2012; Vucic-Pestic et al., 2010). Previous studies on spiders, including another species of jumping spider, have also found lower feeding rates in males and have suggested that these lower feeding rates may be the result of the differences in reproductive roles among the sexes (Givens, 1978; Walker & Rypstra, 2002). These studies suggest that females may forage to maximize energy intake for the development of eggs while males may feed to meet a minimum energy requirement allowing them to devote more time to pursue mating opportunities. Therefore, differences in the fitness benefits of foraging among sexes and stages may shape the differences in functional responses in this system. Furthermore, the difference in reproductive roles is potentially what has allowed for the evolution of sexually dimorphic characteristics such as the enlarged chelicerae of males that may detrimentally affect foraging.

Temperature is widely thought to play an important role in determining predator feeding rates by altering predator space clearance rates and handling times, particularly in ectotherms (DeLong, 2021; Englund et al., 2011; Rall et al., 2012; West & Post, 2016). In general, physiological considerations lead to the expectation that space clearance rates should be increasing or unimodal, concave down functions of temperature, whereas handling times are expected to be decreasing or unimodal, concave up functions of temperature (Burnside et al., 2014; DeLong & Lyon, 2020; Englund et al., 2011; Rall et al., 2012; Uszko et al., 2017). Here, we find that jumping spider handling times agree with the expectation that handling times will decrease monotonically with increasing temperatures. However, we also find little evidence of temperature dependence in the space clearance rates of females and weak evidence for an increase in male and juvenile space clearance rates at high temperatures. Although there is the appearance of no temperature dependence in space clearance rates for females, it is possible that, rather than a unimodal response, the female spiders exhibit an all-or-nothing response to temperature. For example, spiders generally began foraging mid-morning and ceased being present on the walls by mid-afternoon. This is reflected by a unimodal, concave down relationship between female densities and temperature (Supplementary Figure S1.1A). Thus, females may choose to forage only under temperatures in which they forage effectively. This could lead to the lack of temperature dependence in space clearance rates we observed despite a still-possible physiological response to temperature. It is also possible that, as is common in other ectotherms, zebra jumping spiders can regulate their body temperature behaviorally by using temperature refuges and microhabitats on the walls. The potential increase in juvenile and male space clearance rates at high temperatures may also be reflected in the patterns of spider densities with temperature. Males and juveniles do not show as strong of a unimodal relationship with temperature as females, particularly at high temperatures (Supplementary Figure S1.1B,C). Therefore, the lower density of females at high temperatures may free up males and juveniles to forage at high temperatures. Our results on the temperature dependence of space clearance rates contrast with experimental examinations of temperature effects on functional responses in which individuals are often exposed to a uniform temperature and are typically in homogeneous arenas with no potential temperature refuges (Archer et al., 2019; Broom et al., 2021; Islam et al., 2021; Russell et al., 2021) This difference has potentially strong implications for the general way in which we conceive of temperature altering predator-prey interactions and thus food web dynamics given changing climates. Using more realistic temperature regimes and arenas in experiments will allow us to understand whether our result is unique to zebra jumping spiders or potentially more widespread in terrestrial ectotherms.

Mutual interference is also known to be a widespread factor influencing predator feeding rates (DeLong & Vasseur, 2011; Novak & Stouffer, 2021). Most studies on interference measure its effects by altering the densities of similar predators and examining the effects on predator feeding rates. This has limited most previous studies from examining how intraspecific differences might influence interference rates. Our results suggest that interference rates may be asymmetric within species. For example, interference showed little effect on male feeding rates relative to females and juveniles. For females and juveniles, we also found evidence that the strength of interference may depend on the sex/stage from which the interference originates. In particular, females showed stronger interference from males and juveniles. This could be because of the time spent interacting or mating with males and the potential pursuit of juveniles as prey, as cannibalism of juveniles occurs in this system (K.E. Coblentz; *Personal Observation*). Juveniles showed greater interference from the adult stages, which also may be a product of being potential prey of the adult stages. If common, this sort of asymmetric interference could have important consequences for population dynamics, demographics, and stage structure as has been found for cannibalism and size-dependent resource competition (Bassar et al., 2017; de Roos & Persson, 2013).

## Conclusions

Predator functional responses are shaped by a multitude of factors. However, our understanding of how these multiple factors might interact with one another to shape predator functional responses is limited by constraints to experimental approaches for measuring functional responses. Using an observational approach, our results reveal interactions between sexes/stages, temperature, and interference in shaping functional responses and raise the possibility of a lack of temperature-dependence in predator space clearance rates under natural conditions and asymmetric intraspecific interference. Further measurements of predator functional responses under field conditions will allow us to gain a better understanding of the multidimensional nature of predator functional responses and, therefore, a better understanding of predator-prey interaction strengths and their consequences.

## Acknowledgements

We thank Troy Scheer for measuring spiders from photographs taken during the feeding surveys, Lyndsie Wszola, Miranda Salsbery, and Francis Biagioli for feedback on the statistical analyses, Ben DeLong for calculating the areas of the walls, and the staff of Cedar Point Biological Station for making our research possible during a pandemic. Funding for this research was provided by a James S. McDonnel Foundation Scholar Award in Studying Complex Systems to JPD and an NSF Graduate Research Fellowship to SFU (DGE-1610400).

## Author Contributions

KEC and JPD designed the research, all authors performed the feeding surveys, KEC performed the detection time observations, performed the statistical analyses, and wrote the first draft of the manuscript. All authors contributed to revisions.

## Conflict of Interest

The authors declare no conflicts of interest.

## Data Availability Statement

All data and code for their analyses are available on GitHub: https://github.com/KyleCoblentz/ZebraSpiderFR. Upon acceptance, the GitHub repository will be permanently archived on Zenodo.

## Notes

### Competing Interest Statement

The authors have declared no competing interest.

## References

Amarasekare, P. (2015). Effects of temperature on consumer-resource interactions. Journal of Animal Ecology, 84(3), 665–679. https://doi.org/10.1111/1365-2656.12320

Archer, L. C., Sohlström, E. H., Gallo, B., Jochum, M., Woodward, G., Kordas, R. L., Rall, B. C., & O’Gorman, E. J. (2019). Consistent temperature dependence of functional response parameters and their use in predicting population abundance. Journal of Animal Ecology, 88(11), 1670–1683. https://doi.org/10.1111/1365-2656.13060

Bassar, R. D., Travis, J., & Coulson, T. (2017). Predicting coexistence in species with continuous ontogenetic niche shifts and competitive asymmetry. Ecology, 98(11), 2823–2836. https://doi.org/10.1002/ecy.1969

Baudrot, V., Perasso, A., Fritsch, C., Giraudoux, P., & Raoul, F. (2016). The adaptation of generalist predators’ diet in a multi-prey context: Insights from new functional responses. Ecology, 97(7), 1832-1841.

Beardsell, A., Gravel, D., Berteaux, D., Gauthier, G., Clermont, J., Careau, V., Lecomte, N., Juhasz, C.-C., Royer-Boutin, P., & Bêty, J. (2021). Derivation of Predator Functional Responses Using a Mechanistic Approach in a Natural System. Frontiers in Ecology and Evolution, 0. https://doi.org/10.3389/fevo.2021.630944

Beddington, J. R. (1975). Mutual Interference Between Parasites or Predators and its Effect on Searching Efficiency. Journal of Animal Ecology, 44(1), 331–340. https://doi.org/10.2307/3866

Broom, C. J., South, J., & Weyl, O. L. F. (2021). Prey type and temperature influence functional responses of threatened endemic Cape Floristic Ecoregion fishes. Environmental Biology of Fishes, 104(7), 797–810. https://doi.org/10.1007/s10641-021-01111-w

Bürkner, P.-C. (2017). brms: An R Package for Bayesian Multilevel Models Using Stan. Journal of Statistical Software, 80(1), 1–28. https://doi.org/10.18637/jss.v080.i01

Burnside, W. R., Erhardt, E. B., Hammond, S. T., & Brown, J. H. (2014). Rates of biotic interactions scale predictably with temperature despite variation. Oikos, 123(12), 1449–1456. https://doi.org/10.1111/oik.01199

Coblentz, K. E., & DeLong, J. P. (2020). Predator-dependent functional responses alter the coexistence and indirect effects among prey that share a predator. Oikos, 129(9), 1404–1414. https://doi.org/10.1111/oik.07309

de Roos, A. M., & Persson, L. (2013). Population and Community Ecology of Ontogenetic Development. Princeton University Press. https://www.jstor.org/stable/j.ctt1r2g73

DeLong, J. P. (2021). Predator Ecology: Evolutionary Ecology of the Functional Response. Oxford University Press.

DeLong, J. P., & Lyon, S. (2020). Temperature alters the shape of predator-prey cycles through effects on underlying mechanisms. PeerJ, 8, e9377. https://doi.org/10.7717/peerj.9377

DeLong, J. P., & Vasseur, D. A. (2011). Mutual interference is common and mostly intermediate in magnitude. BMC Ecology, 11(1), 1. https://doi.org/10.1186/1472-6785-11-1

Dick, J. T. A., Alexander, M. E., Jeschke, J. M., Ricciardi, A., MacIsaac, H. J., Robinson, T. B., Kumschick, S., Weyl, O. L. F., Dunn, A. M., Hatcher, M. J., Paterson, R. A., Farnsworth, K. D., & Richardson, D. M. (2014). Advancing impact prediction and hypothesis testing in invasion ecology using a comparative functional response approach. Biological Invasions, 16(4), 735–753. https://doi.org/10.1007/s10530-013-0550-8

Englund, G., Öhlund, G., Hein, C. L., & Diehl, S. (2011). Temperature dependence of the functional response. Ecology Letters, 14(9), 914–921. https://doi.org/10.1111/j.1461-0248.2011.01661.x

Gilbert, B., Tunney, T. D., McCann, K. S., DeLong, J. P., Vasseur, D. A., Savage, V., Shurin, J. B., Dell, A. I., Barton, B. T., Harley, C. D. G., Kharouba, H. M., Kratina, P., Blanchard, J. L., Clements, C., Winder, M., Greig, H. S., & O’Connor, M. I. (2014). A bioenergetic framework for the temperature dependence of trophic interactions. Ecology Letters, 17(8), 902–914. https://doi.org/10.1111/ele.12307

Givens, R. P. (1978). Dimorphic Foraging Strategies of a Salticid Spider (Phidippus Audax). Ecology, 59(2), 309–321. https://doi.org/10.2307/1936376

Griffen, B. D. (2021). Considerations When Applying the Consumer Functional Response Measured Under Artificial Conditions. Frontiers in Ecology and Evolution, 0. https://doi.org/10.3389/fevo.2021.713147

Hassell, M. P., & Varley, G. C. (1969). New Inductive Population Model for Insect Parasites and its Bearing on Biological Control. Nature, 223(5211), 1133–1137. https://doi.org/10.1038/2231133a0

Holling, C. S. (1959). Some Characteristics of Simple Types of Predation and Parasitism. The Canadian Entomologist, 91(7), 385–398. https://doi.org/10.4039/Ent91385-7

Hossie, T. J., Chan, K., & Murray, D. L. (2021). Increasing availability of palatable prey induces predator-dependence and increases predation on unpalatable prey. Scientific Reports, 11(1), 6763. https://doi.org/10.1038/s41598-021-86080-x

Islam, Y., Shah, F. M., Rubing, X., Razaq, M., Yabo, M., Xihong, L., & Zhou, X. (2021). Functional response of Harmonia axyridis preying on Acyrthosiphon pisum nymphs: The effect of temperature. Scientific Reports, 11(1), 13565. https://doi.org/10.1038/s41598-021-92954-x

McElreath, R. (2016). Statistical Rethinking: A Bayesian Course with Examples in R and Stan. Chapman and Hall/CRC. https://doi.org/10.1201/9781315372495

Mocq, J., Soukup, P. R., Näslund, J., & Boukal, D. S. (2021). Disentangling the nonlinear effects of habitat complexity on functional responses. Journal of Animal Ecology, 90(6), 1525–1537. https://doi.org/10.1111/1365-2656.13473

Murdoch, W. W. (1969). Switching in General Predators: Experiments on Predator Specificity and Stability of Prey Populations. Ecological Monographs, 39(4), 335–354. https://doi.org/10.2307/1942352

Murdoch, W. W., & Oaten, A. (1975). Predation and Population Stability. In A. MacFadyen (Ed.), Advances in Ecological Research (Vol. 9, pp. 1–131). Academic Press. https://doi.org/10.1016/S0065-2504(08)60288-3

Novak, M., & Stouffer, D. B. (2021). Systematic bias in studies of consumer functional responses. Ecology Letters, 24(3), 580–593. https://doi.org/10.1111/ele.13660

Novak, M., Wolf, C., Coblentz, K. E., & Shepard, I. D. (2017). Quantifying predator dependence in the functional response of generalist predators. Ecology Letters, 20(6), 761–769. https://doi.org/10.1111/ele.12777

Novak, M., & Wootton, J. T. (2008). Estimating Nonlinear Interaction Strengths: An Observation-Based Method for Species-Rich Food Webs. Ecology, 89(8), 2083–2089. https://doi.org/10.1890/08-0033.1

O’Connor, M. I., Gilbert, B., & Brown, C. J. (2011). Theoretical Predictions for How Temperature Affects the Dynamics of Interacting Herbivores and Plants. The American Naturalist, 178(5), 626–638. https://doi.org/10.1086/662171

Otto, S. B., Rall, B. C., & Brose, U. (2007). Allometric degree distributions facilitate food-web stability. Nature, 450(7173), 1226–1229. https://doi.org/10.1038/nature06359

Preston, D. L., Henderson, J. S., Falke, L. P., Segui, L. M., Layden, T. J., & Novak, M. (2018). What drives interaction strengths in complex food webs? A test with feeding rates of a generalist stream predator. Ecology, 99(7), 1591–1601. https://doi.org/10.1002/ecy.2387

R Core Team. (2020). R: A language and environment for statistical computing. R Foundation for Statistical Computing, Vienna, Austria. https://www.R-project.org/

Rall, B. C., Brose, U., Hartvig, M., Kalinkat, G., Schwarzmüller, F., Vucic-Pestic, O., & Petchey, O. L. (2012). Universal temperature and body-mass scaling of feeding rates. Philosophical Transactions of the Royal Society B: Biological Sciences, 367(1605), 2923–2934. https://doi.org/10.1098/rstb.2012.0242

Russell, M. C., Qureshi, A., Wilson, C. G., & Cator, L. J. (2021). Size, not temperature, drives cyclopoid copepod predation of invasive mosquito larvae. PLOS ONE, 16(2), e0246178. https://doi.org/10.1371/journal.pone.0246178

Schneider, C. A., Rasband, W. S., & Eliceiri, K. W. (2012). NIH Image to ImageJ: 25 years of image analysis. Nature Methods, 9(7), 671-675.https://doi.org/10.1038/nmeth.2089

Smout, S., Asseburg, C., Matthiopoulos, J., Fernández, C., Redpath, S., Thirgood, S., & Harwood, J. (2010). The Functional Response of a Generalist Predator. PLOS ONE, 5(5), e10761. https://doi.org/10.1371/journal.pone.0010761

Solomon, M. E. (1949). The Natural Control of Animal Populations. Journal of Animal Ecology, 18(1), 1–35. https://doi.org/10.2307/1578

Thompson, D. J. (1978). Towards a Realistic Predator-Prey Model: The Effect of Temperature on the Functional Response and Life History of Larvae of the Damselfly, Ischnura elegans. Journal of Animal Ecology, 47(3), 757–767. https://doi.org/10.2307/3669

Toscano, B. J., & Griffen, B. D. (2013). Predator size interacts with habitat structure to determine the allometric scaling of the functional response. Oikos, 122(3), 454–462. https://doi.org/10.1111/j.1600-0706.2012.20690.x

Uiterwaal, S. F., & DeLong, J. P. (2020). Functional responses are maximized at intermediate temperatures. Ecology, 101(4), e02975. https://doi.org/10.1002/ecy.2975

Uiterwaal, S. F., Lagerstrom, I. T., Lyon, S. R., & DeLong, J. P. (2018). Data paper: FoRAGE (Functional Responses from Around the Globe in all Ecosystems) database: a compilation of functional responses for consumers and parasitoids [Preprint]. Ecology; bioRxiv. https://doi.org/10.1101/503334

Uszko, W., Diehl, S., Englund, G., & Amarasekare, P. (2017). Effects of warming on predator-prey interactions – a resource-based approach and a theoretical synthesis. Ecology Letters, 20(4), 513–523. https://doi.org/10.1111/ele.12755

Vehtari, A., Gabry, J., Magnusson, M., Yao, Y., Bürkner, P.-C., Paananen, T., Gelman, A., Goodrich, B., Piironen, J., & Nicenboim, B. (2020). loo: Efficient Leave-One-Out Cross-Validation and WAIC for Bayesian Models (2.4.1) [Computer software]. https://CRAN.R-project.org/package=loo

Vucic-Pestic, O., Rall, B. C., Kalinkat, G., & Brose, U. (2010). Allometric functional response model: Body masses constrain interaction strengths. Journal of Animal Ecology, 79(1), 249–256. https://doi.org/10.1111/j.1365-2656.2009.01622.x

Walker, S. E., & Rypstra, A. L. (2002). Sexual dimorphism in trophic morphology and feeding behavior of wolf spiders (Araneae: Lycosidae) as a result of differences in reproductive roles. Canadian Journal of Zoology, 80(4), 679–688. https://doi.org/10.1139/z02-037

Watanabe, S. (2013). A Widely Applicable Bayesian Information Criterion. Journal of Machine Learning Research, 14(Mar), 867-897.

West, D. C., & Post, D. M. (2016). Impacts of warming revealed by linking resource growth rates with consumer functional responses. Journal of Animal Ecology, 85(3), 671–680. https://doi.org/10.1111/1365-2656.12491

