## Supplemental Material S1 for "Quantifying predator functional responses under field conditions reveals interactive effects of temperature and interference with sex and stage"

**Table S1.1.** Functional response models fit for female zebra jumping spiders with Widely Applicable Information Criterion (WAIC) values, differences between the top WAIC score and lower ranked models Δ WAIC, the standard error in those difference SE Δ WAIC, model weights for each functional response model, and the estimated parameter values for each model (see equation 2 in the main text for parameter definitions). Note that space clearance rates $a$ are reported in hours where as the parameters associated with space clearance rates in temperature-dependent models ($c_a$, $b_a$, and $q_a$) are reported in units of days.

| **Females** | | | | | | | | | | | | | |
|---|---|---|---|---|---|---|---|---|---|---|---|---|---|
| Model | WAIC | ΔWAIC | ΔWAIC SE | Model Weight | $c_a$ | $b_a$ | $q_a$ | $a$ | $\gamma_F$ | $\gamma_M$ | $\gamma_J$ | $\gamma$ | $f$ |
| No Temperature Interference Separate | 351.2 | 0 | 0 | 0.43 | -- | -- | -- | 2.63 (2.1, 2.82) | 0.34 (0.01, 1.15) | 1.00 (0.05, 2.73) | 1.05 (0.04, 3.23) | -- | -- |
| No Temperature Interference Combined | 351.5 | 0.3 | 1.6 | 0.37 | -- | -- | -- | 2.42 (1.94, 3.04) | -- | -- | -- | 0.35 (0.02, 0.94) | -- |
| No Temperature No Interference | 353.7 | 2.5 | 3 | 0.12 | -- | -- | -- | 2.1 (1.76, 2.46) | -- | -- | -- | -- | -- |
| + Temperature Interference Combined | 356.0 | 4.8 | 3.6 | 0.04 | 25.02 (6.45, 49.33) | 0.10 (0.016, 0.20) | -0.001 (-0.004, 0.0001) | -- | -- | -- | -- | 1.43 (0.18, 3.74) | -- |
| + Temperature Interference Separate | 357.3 | 6.1 | 4.2 | 0.02 | 25.58 (7.18, 49.15) | 0.10 (0.03, 0.19) | -0.002 (-0.004, -0.00006) | -- | 1.42 (0.08, 4.3) | 2.48 (0.24, 6.23) | 2.25 (0.09, 6.32) | -- | -- |
| + Temperature No Interference | 357.9 | 6.7 | 4 | 0.02 | 25.7 (7.64, 48.4) | 0.06 (-0.003, 0.15) | -0.001 (-0.003,0.0004) | -- | -- | -- | -- | -- | -- |
| Null | 450.4 | 99.2 | 25 | 0 | -- | -- | -- | -- | -- | -- | -- | -- | 0.92 (0.85, 1.01) |

**Table S1.2.** Functional response models fit for male zebra jumping spiders with Widely Applicable Information Criterion (WAIC) values, differences between the top WAIC score and lower ranked models Δ WAIC, the standard error in those difference SE Δ WAIC, model weights for each functional response model, and the estimated parameter values for each model (see equation 2 in the main text for parameter definitions). Note that space clearance rates $a$ are reported in hours where as the parameters associated with space clearance rates in temperature-dependent models ($c_a$, $b_a$, and $q_a$) are reported in units of days.

| **Males** | | | | | | | | | | | | | |
|---|---|---|---|---|---|---|---|---|---|---|---|---|---|
| Model | WAIC | ΔWAIC | ΔWAIC SE | Model Weight | $c_a$ | $b_a$ | $q_a$ | $a$ | $\gamma_M$ | $\gamma_F$ | $\gamma_J$ | $\gamma$ | $f$ |
| No Temperature No Interference | 116.4 | 0 | 0 | 0.57 | -- | -- | -- | 0.13 (0.12, 0.28) | -- | -- | -- | -- | -- |
| + Temperature No Interference | 118.7 | 2.3 | 1.4 | 0.18 | 19.16 (2.17, 44.09) | -0.1 (-0.22, 0.07) | 0.002 (-0.002, 0.005) | -- | -- | -- | -- | -- | -- |
| No Temperature Interference Combined | 119.8 | 3.4 | 2.4 | 0.11 | -- | -- | -- | 0.4 (0.16, 0.96) | -- | -- | -- | 2.23 (0.04, 8.34) | -- |
| + Temperature Interference Combined | 121.3 | 4.9 | 2.6 | 0.05 | 19.35 (2.11, 44.27) | -0.07 (-0.2, 0.1) | 0.0013 (-0.003,0.005) | -- | -- | -- | -- | 1.34 (0.03, 5.75) | -- |
| No Temperature Interference Separate | 121.6 | 5.2 | 3.2 | 0.04 | -- | -- | -- | 0.49 (0.355, 0.92) | 3.52 (0.17, 9.8) | 3.3 (0.14, 9.52) | 2.98 (0.11, 9.31) | -- | -- |
| Null | 122.5 | 6.1 | 7.8 | 0.03 | -- | -- | -- | -- | -- | -- | -- | -- | 0.18 (0.12, 0.25) |
| + Temperature Interference Separate | 123.9 | 7.5 | 3.6 | 0.02 | 19.2 (2.11, 42.5) | -0.05 (-0.18, 0.12) | 0.001 (-0.003, 0.004) | -- | 2.85 (0.09, 8.88) | 2.44 (0.08, 8.27) | 2.65 (0.08, 8.67) | -- | -- |

**Table S1.3.** Functional response models fit for juvenile zebra jumping spiders with Widely Applicable Information Criterion (WAIC) values, differences between the top WAIC score and lower ranked models Δ WAIC, the standard error in those difference SE Δ WAIC, model weights for each functional response model, and the estimated parameter values for each model (see equation 2 in the main text for parameter definitions). Note that space clearance rates $a$ are reported in hours where as the parameters associated with space clearance rates in temperature-dependent models ($c_a$, $b_a$, and $q_a$) are reported in units of days.

| Juveniles | | | | | | | | | | | | | |
|---|---|---|---|---|---|---|---|---|---|---|---|---|---|
| Model | WAIC | ΔWAIC | ΔWAIC SE | Model Weight | $c_a$ | $b_a$ | $q_a$ | $a$ | $\gamma_J$ | $\gamma_F$ | $\gamma_M$ | $\gamma$ | $f$ |
| No Temperature Interference Separate | 182.7 | 0 | 0 | 0.23 | -- | -- | -- | 1.81 (1.17, 2.57) | 1.208 (0.03, 4.27) | 2.04 (0.15, 5.21) | 1.81 (0.05, 5.6) | -- | -- |
| No Temperature Interference Combined | 182.8 | 0.1 | 1.4 | 0.22 | -- | -- | -- | 1.57 (0.99, 2.33) | -- | -- | -- | 1.03 (0.06, 2.9) | -- |
| + Temperature Interference Separate | 182.8 | 0.1 | 5.2 | 0.21 | 20.84 (3.35, 44.39) | 0.06 (-0.04, 0.18) | -0.0001 (-0.003, 0.002) | -- | 2.09 (0.05, 7.2) | 5.29 (0.83, 11.96) | 3.7 (0.26, 9.83) | -- | -- |
| + Temperature Interference Combined | 183.6 | 0.9 | 4.6 | 0.15 | 20.74 (3.3, 45.16) | 0.04 (-0.06, 0.17) | 0.0001 (-0.002, 0.002) | -- | -- | -- | -- | 3.48 (0.31, 4.72) | -- |
| No Temperature No Interference | 184.1 | 1.4 | 2 | 0.11 | -- | -- | -- | 1.15 (0.82, 1.55) | -- | -- | -- | -- | -- |
| + Temperature No Interference | 184.8 | 2.1 | 4.4 | 0.08 | 20.53 (3.17, 43.96) | -0.01 (-0.09, 0.1) | 0.0009 (-0.002, 0.003) | -- | -- | -- | -- | -- | -- |
| Null | 196.5 | 13.8 | 13 | 0 | -- | -- | -- | -- | -- | -- | -- | -- | |

**Figure S1.1.** Densities of female zebra jumping spiders show a unimodal relationship with temperature (A). The relationship between male and juvenile zebra jumping spider densities with temperature show less of a reduction at high temperatures compared to females (B,C). The lines in the panels represent the fit of a quadratic model between temperature and sex/stage density.

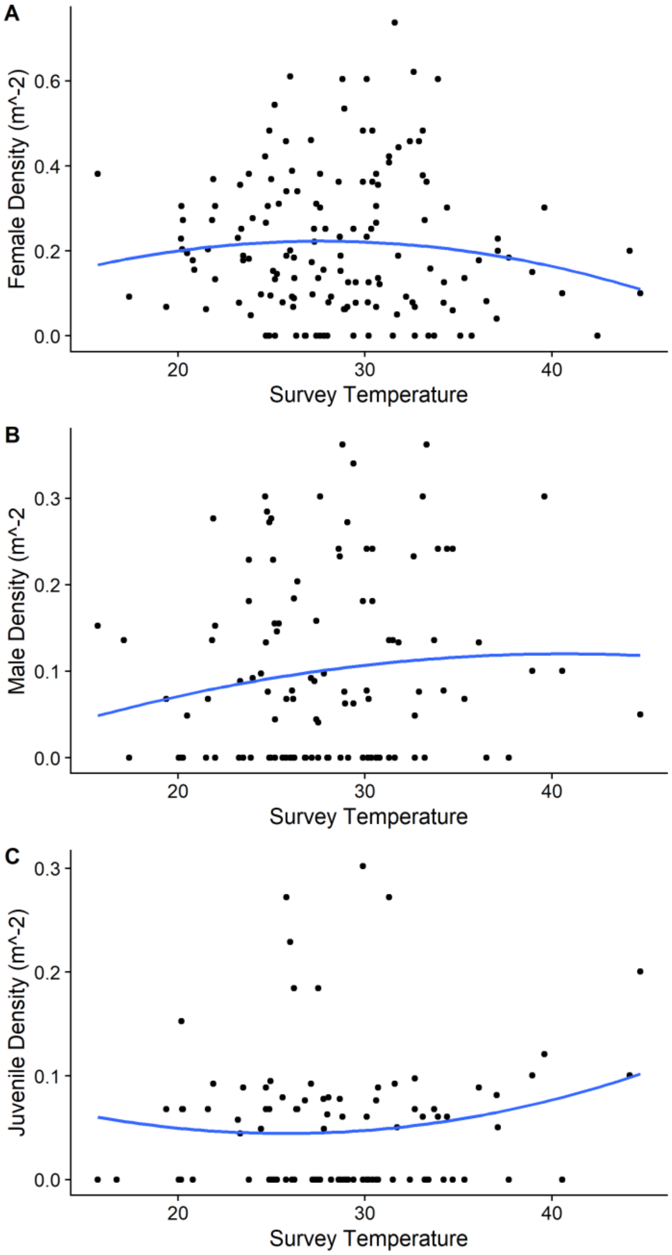
